# Strategic allocation of working memory resource

**DOI:** 10.1101/329870

**Authors:** Aspen H. Yoo, Zuzanna Klyszejko, Clayton E. Curtis, Wei Ji Ma

## Abstract

Visual working memory (VWM), the brief retention of past visual information, supports a range of cognitive functions (Fukuda, Vogel, Mayr, & Awh, 2010; Johnson et al., 2013). The resource that supports VWM is limited, raising the question of how the brain allocates this limited resource to different objects. This question is even more interesting in ecological settings, in which objects are not equally important. In a psychophysical experiment, participants remembered the location of four targets with different probabilities of being tested after a delay. We then measured their memory accuracy of one of the targets. We found that participants allocated more resource to memoranda with higher priority, but underallocated resource to high- and overallocated to low-priority targets relative to the true probe probabilities. These results are well explained by a computational model in which resource is allocated to minimize expected estimation error. We replicated this finding in a second experiment in which participants bet on their memory fidelity after making the location estimate. The results of this experiment show that people 1) use information about memory quality and 2) minimize error even with an incentivized, alternative resource allocation strategy. Humans may mitigate the behavioral effects of a limited VWM through knowledge of memory fidelity and strategic resource allocation.

---

One of the hallmarks of VWM is that it is supported by a limited resource. In natural environments, where objects vary in how relevant they are, the process by which our memory resource is allocated appears flexible and strategic. Indeed, experiments demonstrate that increasing the behavioral relevance of a set of items results in better memory for those items (Bays, 2014; Dube, Emrich, & Al-Aidroos, 2017; Emrich, Lockhart, & Al-Aidroos, 2017; Klyszejko, Rahmati, & Curtis, 2014; Zhang & Luck, 2008). Yet, it is still unknown how people decide how much resource to allocate to the encoding and storing of items with different behavioral relevancies.

Here, our overall objective is to use computational models of VWM performance to understand the strategy by which memory resource is flexibly allocated when items vary in behavioral relevance. To do so, we first established that the amount of allocated resource is monotonically related to the behavioral relevance, or priority, of memorized items. We used a memory-guided saccade task in which, on each trial, participants remembered the location of four dots, one in each visual quadrant (**Fig. 1a**). To operationalize resource prioritization, we used a precue to indicate the probability that each dot would be later probed. On each trial, the probe probabilities were 0.6 (“high”), 0.3 (“medium”), 0.1 (“low”), and 0.0. After a variable delay period, one of the quadrants was cued and the participant made a saccade to the remembered location of the dot within that quadrant. In line with our hypothesis, error decreased monotonically with increasing priority (*F*(1,13) = 13.9, *p*< 0.003), reflecting the intuition that people allocate more resource to a more behaviorally relevant target (**Fig. 1b**).

**Fig. 1.**
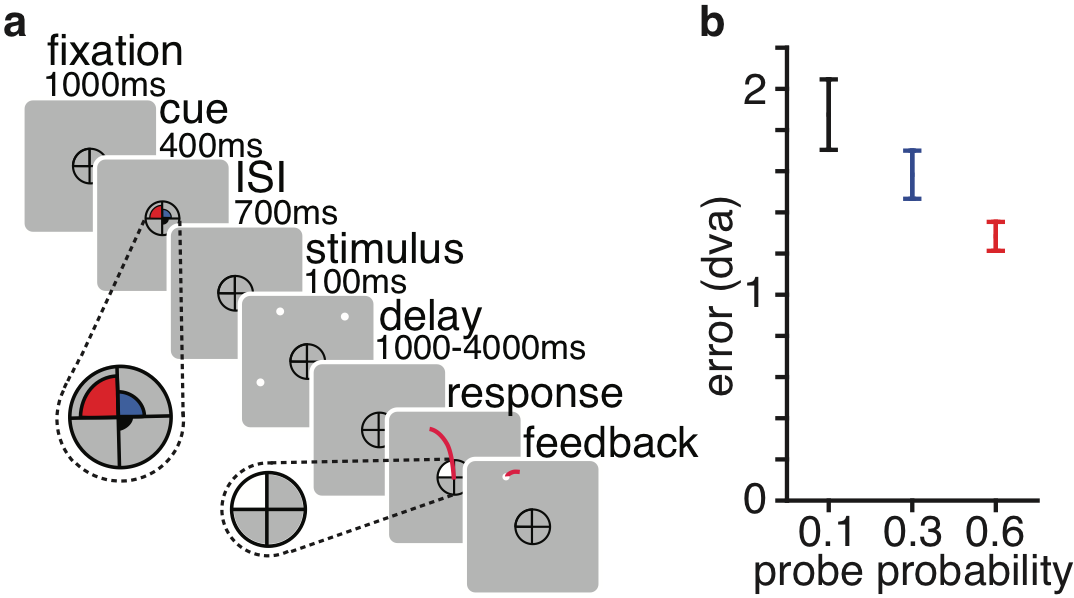
Exp. 1 task sequence and behavioral results. **a**, Task sequence. **b**, Main behavioral effects. Saccade error (*M* ± *SEM*) decreases as a function of increasing priority. red: 0.6., blue: 0.3, black: 0.1.

Next, we asked what strategy people use to allocate resource in response to unequal relevance. Emrich et al. (2017) proposed that resource is allocated in approximate proportion to the probe probabilities. Bays (2014) proposed that resource is allocated such that the expected squared error is minimized. Sims (2015) proposed more generally that resource is allocated to minimize loss. Thus, our second goal was to compare computational models of resource allocation. We tested variable-precision models of estimation errors (Bays & Husain, 2008; van den Berg, Shin, Chou, George, & Ma, 2012) augmented with different resource allocation strategies. Memory precision for a given item is a random variable whose mean depends on the item’s priority (middle/bottom panel of **Fig. 2a**; see Supplementary for more a detailed description of the model). In the *Proportional* model (Emrich et al., 2017), the amount allocated to an item is proportional to the item’s probe probability. This model provided a poor fit to the data (left panel of **Fig. 2b**), suggesting that people do not allocate resource in proportion to probe probability.

Perhaps this model was too constrained, so we tested the *Flexible* model, in which the proportions allocated to each priority condition were free parameters. This model fit the data well (middle panel of **Fig. 2b**) and formal model comparison showed that it outperformed the Proportional model by a median ΔAICc of 63 (bootstrapped 95% CI: [37, 107] and a median ΔBIC of 54 [29, 99]). The proportions allocated to the high-, medium-, and low-priority targets were estimated as 0.49 ± 0.04 (*M* ± *SEM*), 0.28 ± 0.02, and 0.23 ± 0.03, respectively (**Fig. 2c**), suggesting that the brain underallocates resource to high-priority targets and overallocates resource to low-priority targets, relative to the experimental probe probabilities.

The Flexible model offered a good explanation for *how much* participants were allocating to each item, but not *why*. We considered that resource was allocated to minimize expected loss, where loss is defined as estimation error to a power (Bays, 2014; Harris & Wolpert, 1998; Kahneman & Tversky, 1979; Sims, 2015). In this *Minimizing Error* model, resource allocation differs substantially from the Proportional model (De Silva & Ma, 2017). An observer with limited resource should allocate their resource more equally than proportional. Such a strategy would lower the probability of very large errors for low-priority targets, at a small expense of the high-priority targets (top panel of **Fig. 2a**). The exponent on the error serves as a “sensitivity to error” parameter: an observer with a large exponent will experience a large error as much more costly than an observer with a lower exponent, and will adjust their strategy accordingly to avoid those errors. The Minimizing Error model fits better than the Proportional model (median ΔAICc [bootstrapped 95% CI]: 49 [21, 99], ΔBIC: 44 [17, 94]. **Fig. 2b, 2d**) and comparably to the Flexible model (ΔAICc: −7 [−30, −1], ΔBIC: −3 [−26, 3]). Additionally, the model estimated an allocation of resource similar to the Flexible model (0.46 ± 0.02, 0.32 ± 0.01, and 0.22 ± 0.02 for high-, medium-, and low-priority targets, respectively). This suggests that the under- and over-allocation of resources relative to probe probabilities may be rational, stemming from an attempt to minimize error across the experiment.

**Fig. 2.**
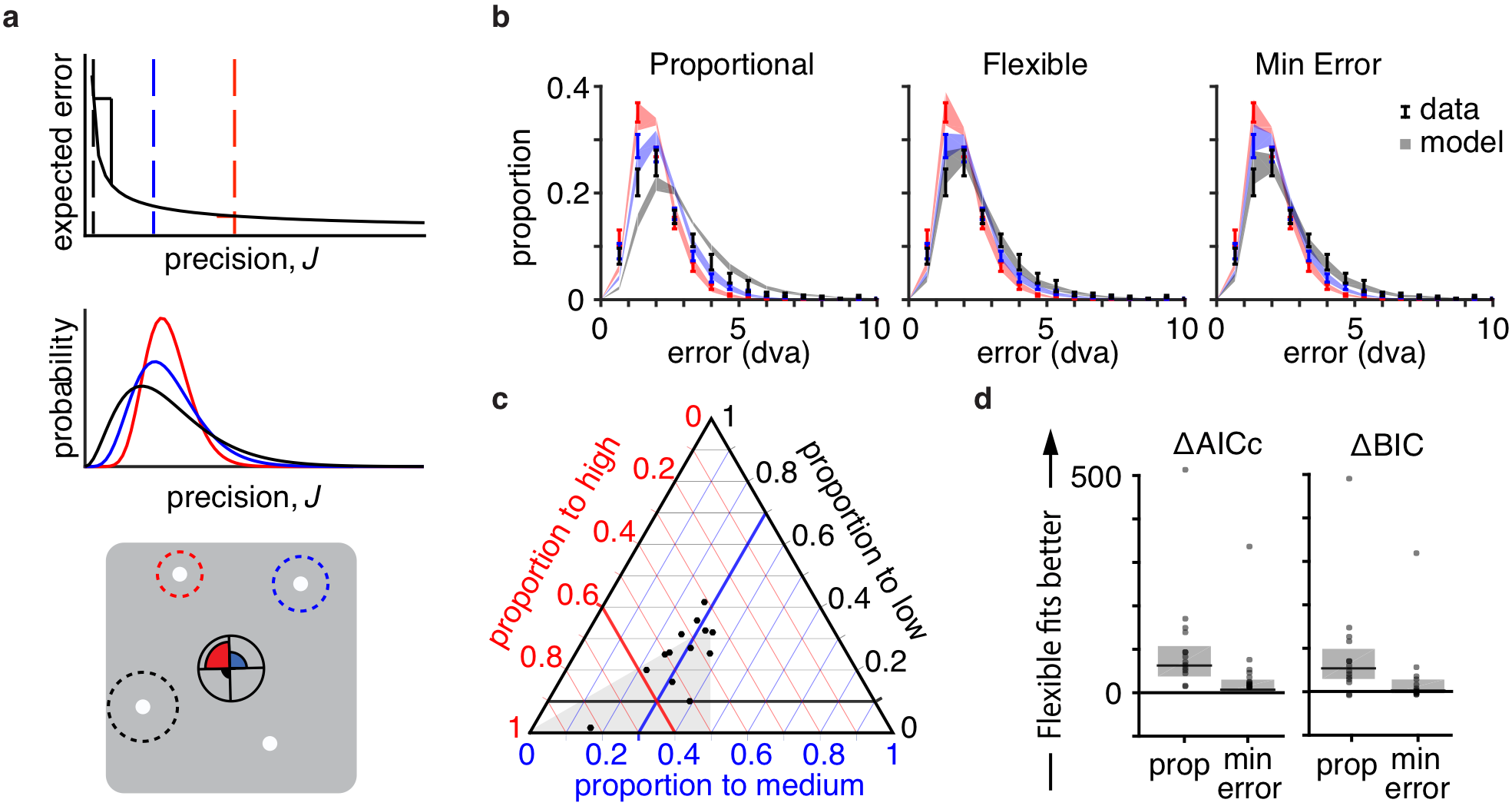
Exp. 1 modeling. Color indicates priority condition – red: 0.6, blue: 0.3, black: 0.1. **a**, Schematic of the Minimizing Error Variable Precision model. *Top*, the expected error of a memory decreases nonlinearly with mean precision. The current amount allocated to each priority item is indicated in the dashed vertical line. An observer can drastically decrease their total expected error by allocating some resource from the high-priority item to the low-priority item. *Middle*, the precision *J* of each item on each trial is drawn from a gamma distribution. Items from different conditions are drawn from distributions with different mean 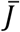, illustrated here in different colors. *Bottom*, the reported location is drawn from a two-dimensional Gaussian with precision *J*; standard deviation 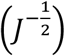 shown in dotted lines. **b**, *M* ± *SEM* error distributions for data (error bars) and model predictions (shaded region) for the Proportional, Flexible, and Minimizing Error models (*N* = 14). **c**, Proportion allocated to each priority condition as estimated from the Flexible model. Black dots represent participants. Thicker lines indicate the 0.6, 0.3, and 0.1 allocation to high, medium, and low, respectively. The intersection of these lines is the prediction for the Proportional model. Observers are underallocating to high priority and overallocating to low, relative to the actual probe probabilities. **d**, Model comparison results. black line: median, grey box: 95% bootstrapped median CI. The Flexible model fits significantly better than the Proportional model, but not significantly better than the Minimizing Error (ME) model.

The first experiment showed that prioritizing items affects memory representations, and that people allocate memory resource in an error-minimizing way. However, this experiment, along with much of the VWM literature, overlooks other information available in VWM: memory uncertainty. Indeed, people can successfully report on the quality of their memory-based decisions (Rademaker, Tredway, & Tong, 2012), suggesting a representation and use of uncertainty over the memorized stimulus (Fougnie, Suchow, & Alvarez, 2011; Honig, Ma, & Fougnie, 2018; van den Berg, Yoo, & Ma, 2017). We conducted a second experiment to investigate how, if at all, priority affects working memory uncertainty.

We tested this with a very similar memory-guided saccade task with an addition wager to measure uncertainty. After the participant made a saccade, a circle appeared centered at the endpoint of the saccade (Graf, Warren, & Maloney, 2005) (**Fig. 3a**). Participants made a wager by adjusting the size of the circle with the goal of enclosing the true target location within the circle. If successful, they received points based on the size of the circle, such that a smaller circle corresponded to more points. In unsuccessful, they received no points. This procedure served as a measure of memory uncertainty because participants were incentivized to make smaller circles when their memory was more certain.

Our predictions for this experiment were the following: a) estimation error decreases with increasing priority, b) circle size decreases with increasing priority, and c) estimation error correlates positively with circle size within each priority level. We confirmed all three predictions. First, estimation error decreased monotonically with increasing priority (*F*(2,20)=12.5, *p*<0.001, *η*^2^=0.55; left panel of **Fig. 3b**), indicating that participants allocated more resource to higher priority targets. Second, circle size decreased monotonically with increasing priority (*F*(1.3,12.9)=10.60, *p*<0.005, *η*^2^=0.51; middle panel of **Fig. 3b**), indicating that participants had higher memory certainty in higher priority trials. Third, estimation error and circle size were correlated within each priority level (*r*_0.6_ = 0.27, *p* < .001; *r*_0.3_ = 0.3, *p* < .001; *r*_0.1_ = 0.2, *p* < .001; right panel of **Fig. 3b**), indicating that people have a single-trial representation of their uncertainty independent of the priority manipulation, as suggested by earlier work (Fougnie et al., 2011; Suchow, Fougnie, & Alvarez, 2017).

The correlation, however, could have been driven by targets closer to the cardinal axes being remembered more precisely then those presented more obliquely (Appelle, 1972; Furmanski & Engel, 2000; Girshick, Landy, & Simoncelli, 2011; Pratte, Park, Rademaker, & Tong, 2017). Pratte and others (2017) argued that much of the seemingly random variability in the variable-precision model could be explained by this “oblique effect.” Perhaps location-dependent noise (along with the observer’s knowledge thereof) might also be driving the measured within-priority correlation. To test this hypothesis, we conducted a permutation test for each participant and priority level (details in Supplementary). We found that the actual correlations (*M* ± *SEM*: 0.29 ± 0.04) were significantly higher than the median of the correlations obtained in the null distribution (*M* ± *SEM*: −0.007 ± 0.006; Wilcoxon signed-rank test, *z* = −4.69, *p* < 1e-5), suggesting that the correlation within each priority condition was driven by internal fluctuations in the quality of the memory representation above and beyond any location-dependent variation.

We extended the computational models from the first experiment to account for the additional wager data. The observer uses trial-to-trial knowledge of memory quality to calculate the probability that the target lies within the circle of a proposed size (a “hit”). The observer multiplies this value by the utility of that circle size to calculate the expected utility (**Fig. 3c**). We assume the observer noisily chooses the circle size that maximizes the expected utility (**Fig. 3d**; details in Supplementary).

**Fig. 3.**
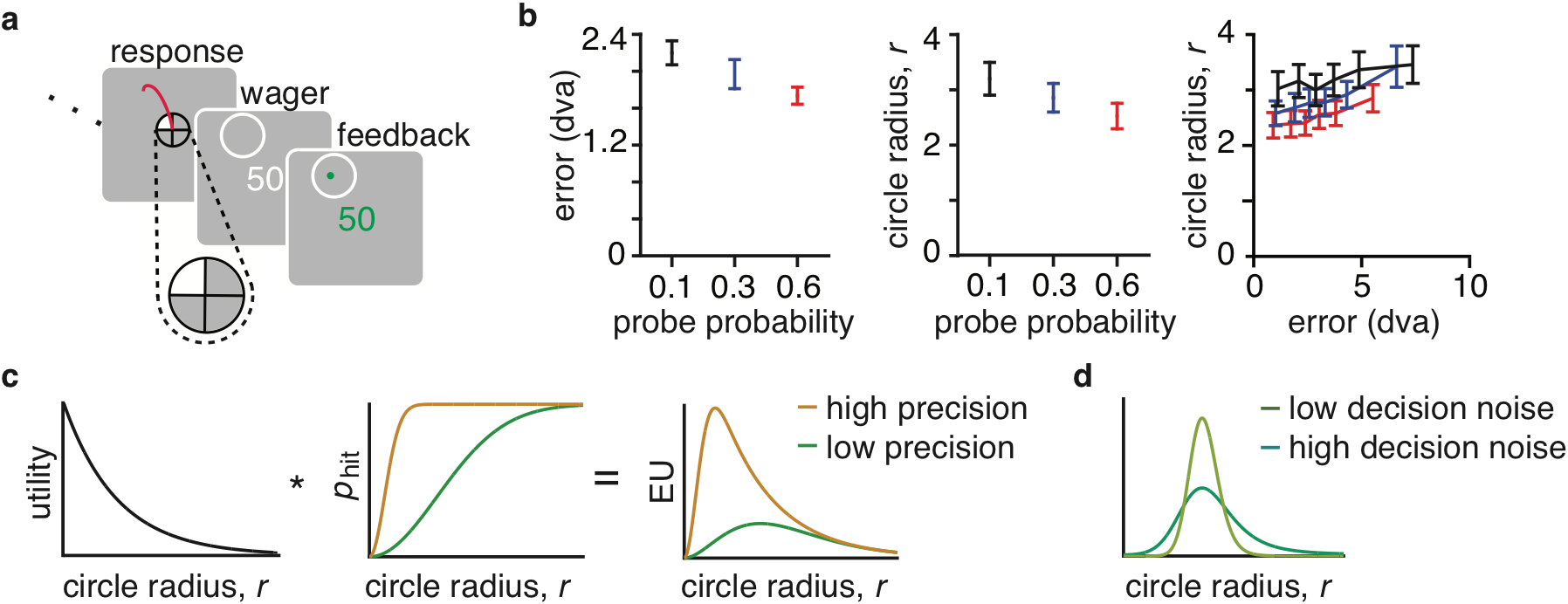
Exp. 2 task, behavior, and model extension. **a**, Trial sequence. Exp. 2 is identical to Exp. 1 up to the saccade response, after which they make the post-decision wager. **b**, Main experimental effects. Error bars show *M* ± *SEM* for memory error (*left*) and circle radius (*middle*) across priorities for 11 participants; both measures decrease with increasing priority. These measures are positively correlated within priority conditions (*right*), suggesting that error and circle size have a common cause, namely fluctuations in precision. **c**, Schematic of how model generates circle radius predictions. For a given radius *r*, the observer multiplies the utility and the probability of the true target being inside of the circle to calculate the expected utility (EU). Shown here are two examples of how precision *J* effects EU (*τ*= 0.1. low precision: *J* = 0.1. high precision: *J* = 2). **d**, To incorporate decision noise, we model response distribution as a softmax function of utility. (low noise: *β* = 1. high noise: *β* = 0.3).

We again tested the Proportional model and Flexible model, jointly fitting the estimation data and the post-estimation wager data. We again found that the Proportional model did not provide a good fit to human data and the Flexible model provided an excellent fit to the data (**Fig. 4a**). As before, the Flexible model suggests that the brain underallocates resource to high-priority targets and overallocates resource to low-priority targets relative to experimental probe probabilities. The proportion allocated to the high-, medium-, and low-priority targets were estimated as 0.44 ± 0.02, 0.31 ± 0.02, and 0.25 ± 0.02, respectively (**Fig. 4b**). Given the analogous results, we again asked if we could describe a normative model for the resource allocation strategy.

Unlike in the first experiment, optimal performance in this experiment requires maximizing points. This *Maximizing Points* model has qualitatively different properties from the Minimizing Error model. An observer that maximizes points would receive more points by ignoring the low-priority targets completely in order to remember the high-priority targets better, while an observer that minimizes error would allocate it more evenly across targets. Because these two strategies conflict, we are able to test whether the intrinsically-driven, error-minimizing strategy that people seem to be using in the absence of reward can withstand being put in conflict with an external incentive. To our surprise, the Maximizing Points model fit very poorly, indicating that participants were not allocating resource in order to earn the most points (Proportional model: median ΔAICc: −75 [−109, −26], ΔBIC: −75 [−109, −26]; Flexible model: ΔAICc: −156 [−308, −94], ΔBIC: −148 [−300, −86]).

Perhaps participants were still acting in in accordance with the Minimizing Error model. In this experiment, this strategy is myopic: the observer allocates resource to minimize error in the estimation without considering how this allocation may affect the points of the wager. Nonetheless, the Minimizing Error model fit the data substantially better than the Proportional model (median ΔAICc: 55 [20, 106], ΔBIC: 50 [17, 102]) and the Maximizing Points model (ΔAICc: 140 [85, 249], ΔBIC: 137 [81, 245]) and about as well as the Flexible model (ΔAICc:−16 [−44, −5], ΔBIC: −12 [−40, 0]; **Fig. 4c**). The Minimizing Error model fitted the proportions of resource allocated to high-, medium-, and low-priority targets as 0.52 ± 0.02, 0.32 ± 0.01, and 0.16 ± 0.01, respectively, similar to the allocation estimated in the Flexible model.

**Fig. 4.**
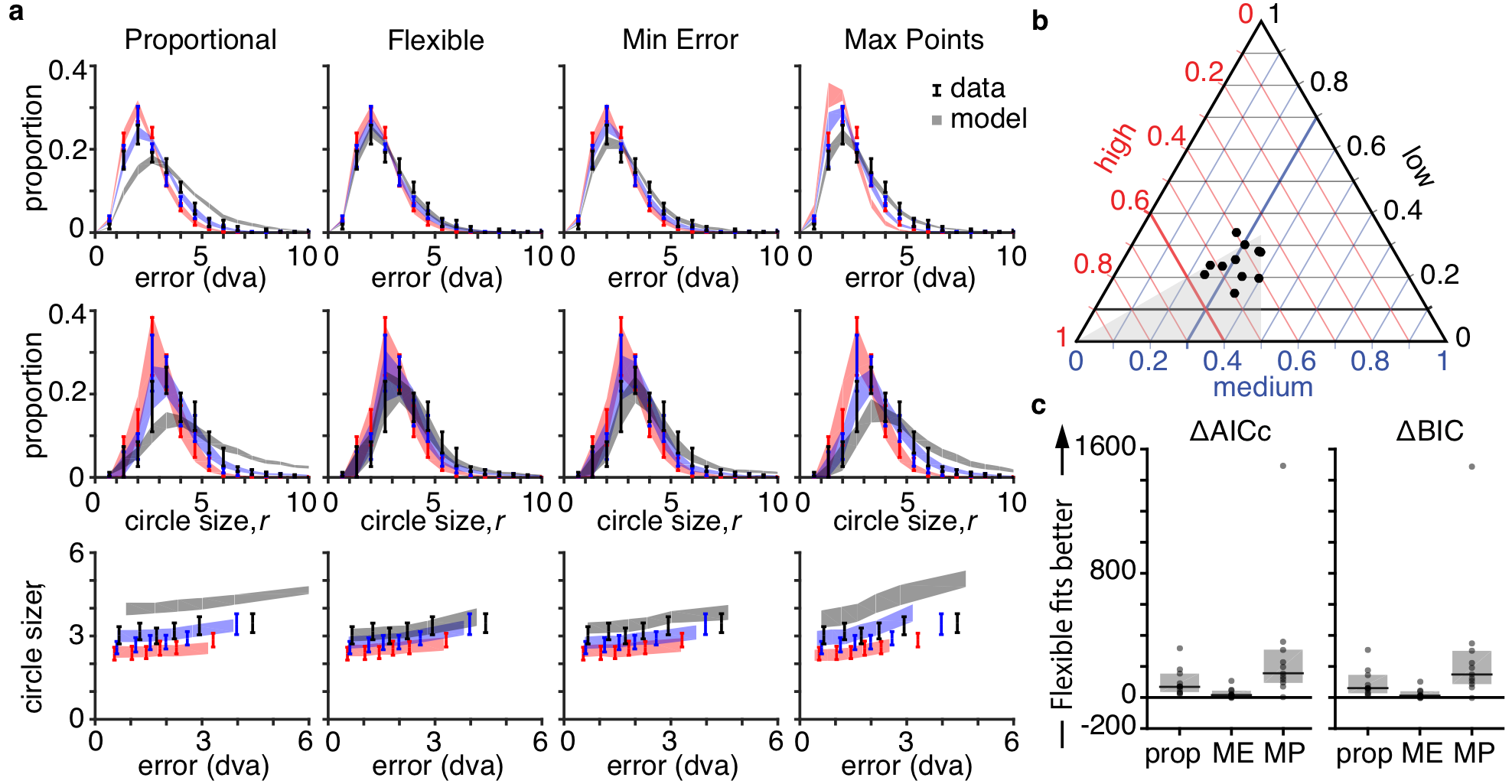
Exp. 2 modeling results. Color indicates priority condition – red: 0.6, blue: 0.3, black: 0.1. **a**, Fits of four models (columns) to error distribution (*top*), circle radius distribution (*middle*), and correlation between the two (*bottom*). *M* ± *SEM* shown for data (error bars) and model predictions (shaded region). **b**, Proportion allocated to each priority condition as estimated from the Flexible model. Black dots represent participants. Thicker lines indicate the 0.6, 0.3, and 0.1 allocation to high, medium, and low, respectively. The intersection of these lines is the prediction for the Proportional model. Again, observers are underallocating to high priority and overallocating to low, relative to the actual probe probabilities. **c**, Model comparison results. black line: median, grey box: 95% bootstrapped median CI. The Flexible model fits significantly better than the Proportional and Maximizing Points (MP) models, but not significantly better than the Minimizing Error (ME) model.

In this work, we examined how people allocate resource in a task with varying behavioral relevance. First, we found that people flexibly allocate resource according to behavioral relevance. This is comforting: we remember more important things better. Second, we found accurate knowledge of both priority-driven and spontaneous trial-to-trial fluctuations in memory quality, even when controlling for spatial location. This is also comforting: to some extent, we can trust our confidence in our memories. Additionally, uncertainty is useful when deciding whether to use or resample information. Third, we explained not just how people allocate VWM resource, but what strategy they may be using. We find that people minimize estimation error, a strategy whose consequence is an overallocation to low- and underallocation to high-priority targets relative to probe probabilities. Fourth, this strategy persists even when presented with a conflicting external reward.

Consider that we find minimizing the errors of our memory intrinsically rewarding. Indeed, extrinsic rewards influence the metrics of saccades in humans and monkeys (Chen, Chen, Zhou, & Mustain, 2014; Takikawa, Kawagoe, Itoh, Nakahara, & Hikosaka, 2002). For instance, extrinsic rewards affect both the velocity of saccades as well as neural activity in dopamine-associated reward circuits (Kato et al., 1995), and they modulate neural activity in cortical areas that represent the goals of saccade plans (Platt & Glimcher, 1999). Perhaps the intrinsic reward associated with veridical memory eclipses the extrinsic reward associated with gaining more points, which would explain why people minimized error instead of maximized points in the second experiment. Additionally, minimizing memory error might be computationally easier than maximizing points because it does not require the observer to think and optimize performance two steps ahead. Lastly, the amount that performance may improve from maximizing points may not be worth the computational and metabolic cost.

Our results identify a single and simple model of how the resource that supports VWM is allocated despite the large variability in WM abilities across individuals and ages (Engle, Kane, & Tuholski, 1999; Salthouse, Babcock, & Shaw, 1991). For example, electrophysiological signals measured at the scalp predict individual differences in WM capacity (Vogel & Machizawa, 2004) as well as trial-to-trial variation in the precision of WM (Adam, Robison, & Vogel, 2018; Reinhart et al., 2012). Individual differences in WM can also be explained by differences in control processes, such as inhibition of irrelevant distractors (Vogel, McCollough, & Machizawa, 2005). Our model accounts for these individual differences by explicitly assuming inter-trial variability, as well as having parameters that account for participants’ differences in total amount of resource as well as sensitivity to error. Some participants prefer making a few large errors in order to maximize the number of extremely precise memory guided saccades, while others prefer avoiding large errors at the expense of those precise saccades.

In everyday life, we are bombarded with information constantly, and we have to decide what to look at, pay attention to, and remember. This study finds that not only do people remember more important items and their associated uncertainty, but they also do so in a way that minimizes the overall magnitude of memory errors. Perhaps this strategic allocation is how we are able to function so well despite such limited working memory resource.

## Methods

### Participants

Fourteen participants (5 males, mean age=30.3, SD=7.2) participated in Experiment 1 and eleven (5 males, age=28.6, SD=3.03) in Experiment 2. Everyone had normal or corrected-to-normal vision and no history of neurological disorders. Participants were naive to the study hypotheses and were paid $10/hour. All participants provided written consent. The study conformed to the Declaration of Helsinki and was approved by the Institutional Review Board of New York University.

### Apparatus

Participants were placed 56 cms from the monitor (19 inches, 60 Hz), with their heads in a chinrest. Eye movements were callibrated using the 9-point calibration and recorded at a frequency of 1000 Hz (Eyelink 1000, SR Research). Target stimuli were programmed in MATLAB (MathWorks) using the MGL toolbox (Gardner Lab, Stanford) and were displayed against a uniform grey background.

In Experiment 2, participants made behavioral responses using a space bar with their left hand and a circular knob (PowerMate, Griffin Technology) with their right hand. For eye-tracking, we applied an online drift correction when the recorded location of center of fixation exceeded 1 degree of visual angle from the center of the fixation cross. Because this experiment provided live visual feedback of the participants’ current fixation, large discrepancies were uncomfortable for the participant and resulted in imprecise data. This was not a problem for the Experiment 1 because the corrective saccade provided a measure of drift, which we used to correct offline.

### Trial procedure

Each trial (**Fig. 1a**) began with a 300 ms increase in the size of the fixation symbol, an encircled fixation cross. This was followed by a 400 ms endogenous precue, consisting of three colored wedges presented within the fixation symbol, each of which angularly filled one quadrant. The radial sizes and colors (pink, yellow, and blue) of the wedges corresponded to probe probabilities of 0.6, 0.3, and 0.1, respectively. The quadrant with a probe probability of 0.0 did not have a wedge.

The precue was presented for 400 ms; this was followed by a 700 ms interstimulus interval, then by the targets, presented for 100 ms. The targets were four dots, each in separate visual quadrants. The dots were presented at approximately 10 dva from fixation, with random jitter of 1 degree of visual angle to each location. The location of the targets in polar coordinates were pseudo-randomly sampled from every 10 degrees, avoiding cardinal axes.

This was followed by a variable delay, chosen with equal probability from the range between 1000 and 4000 ms in 500 ms increments. A response cue appeared afterward, which was a white wedge that filled an entire quadrant of the fixation symbol. Participants were instructed to saccade to the remembered dot location within the corresponding quadrant of the screen. If participants took shorter than 100 ms or longer than 1200 ms to make the saccade, the trial was discarded.

In Experiment 1, after the saccade, the actual dot location was presented as feedback and the participant made a corrective saccade to that location. After 500 ms, the feedback disappeared, participants returned their gaze to the central fixation cross, and a 1500 ms intertrial interval began.

In Experiment 2, simultaneously with the response cue, a red dot appeared at the location of the participants’ fixation as measured online by the eye tracker. Because of eyetracker noise, the red dot often appeared in a slightly different location than where the participant was fixating. Thus, participants were instructed to adjust their gaze such that the red dot was at the remembered location, and press the space bar to indicate that this was their intended saccade endpoint. After completing this response, participants performed a post-estimation wager. A circle appeared, centered at the saccade endpoint. Participants received points based on the size of the circle, such that a smaller circle corresponded to more points. However, participants were only rewarded points if the true target was within the circle. The number of points awarded was 120*e*^−0.4*r*^, in which *r* was the radius of the circle.

### Data Processing

Processing and manual scoring of eye movement data was performed in an inhouse MATLAB function-graphing toolbox (iEye). Eye position and saccadic reaction time (SRT) were extracted from iEye. Statistical analyses were performed in MATLAB (Mathworks) and SPSS (IBM). We excluded trials in which a) participants were not fixating in the middle of the screen during stimulus presentation, b) saccades were initiated before 100 ms or after 1200 ms after the response cue onset, c) pupil data during the response period were missing, or d) participants made a saccade to the wrong quadrant, ignoring the response cue. This resulted in removing between 1% and 7% of trials per subject. Raw gaze positions were transformed offline into degrees of visual angle using a third order polynomial algorithm that fit eye positions to known spatial locations and we used gaze velocity trace to determine the onset and offset of saccades with a $30°/s threshold. Priority effects were not significantly different between initial and final saccade position, so we report the results for the final saccades.

### Model fitting, Parameter Recovery, and Model Recovery

For each participant and each model, we estimated the parameters using maximum-likelihood estimation. The likelihood of the parameters are defined as *p*(data|model,θ), in which θ is a vector of the model parameters. To calculate the parameter likelihood, we use numerical integration to marginalize over the internal variables **x** and *J* (Supplementary Information). To find the maximum-likelihood parameter estimate, we used the optimization algorithm Bayesian Adaptive Direct Search (Acerbi & Ma, 2017) in MATLAB, which combines mesh grid and Bayesian optimization methods. We completed 50 optimizations with different starting values for each participant and model, to ensure the obtained estimates were not a result of a local minimum. We took the maximum of all the runs as our estimate of the maximum-likelihood, and the corresponding parameter combination as our ML parameter estimates.

To validate the data-generating and model-fitting code, we performed parameter and model recovery. We simulated data from each model then fit each model to the simulated data. Successful parameter recovery occurs when the estimated parameters for the model that generated the data are equivalent or close to the true parameters. Parameter recovery is necessary for the interpretability of the parameter estimates. Successful model recovery occurs when the model which generated the data also fits the data better than any other model. Model recovery is necessary to ensure the models are distinguishable in a psychologically plausible model space. We successfully recovered both the parameters and models for each model.

### Data Availability

The datasets generated and analyzed during the current study are available in the WM_resource_allocation github repository: github.com/aspenyoo/WM_resource_allocation.

### Code Availability

All analyses were conducted in MATLAB. Code used to fit and compare models and generate figures are available on the WM_resource_allocation github repository: github.com/aspenyoo/WM_resource_allocation.

## Contributions

All authors designed experiment, ZK collected data, ZK and AHY analyzed data, all authors wrote manuscript.

## Acknowledgements

We thank Maija Honig and Daryl Fougnie for methodological and statistical inspiration and discussion and Helena Palmieri for assisting with data collection. This work was supported by NIH R01-EY016407, NIH R01-EY027925, and the NEI Visual Neuroscience Training Program (T32 EY007136).

